# Spectral Correlates of Encoding Distinguish Good from Poor Learners

**DOI:** 10.64898/2026.07.29.741612

**Authors:** Ben Falkenburg, Adam Broitman, Michael J. Kahana

## Abstract

Individuals vary in their ability to encode and recall new experiences, yet neural markers of memory have rarely been linked to these trait-level differences. Subsequent memory effects (SMEs) identify patterns of neural activity during encoding that predict whether an item will later be remembered or forgotten. In scalp EEG, SMEs are typically characterized by increased theta and gamma power with decreased alpha, a profile that has been interpreted as a signature of effective encoding. Here, we tested whether SMEs reflect individual differences in mnemonic ability by analyzing over 1.3 million encoding events from a large free recall dataset. At the group level, alpha suppression and theta enhancement predicted successful encoding, while gamma did not predict subsequent memory after controlling for serial position. Across individuals, better learners showed reduced SMEs in all three frequency bands. The neural contrast that defines successful encoding was therefore most pronounced among the poorest learners. These findings suggest that high-performing individuals are less reliant on phasic spectral activity for successful encoding, and that SMEs capture stable individual differences in encoding efficiency.

**Highlights:**

- EEG activity during learning differentiates better from poorer learners
- EEG markers of successful encoding also reflect stable learning ability
- Higher-performing individuals show attenuated theta–alpha–gamma signatures
- EEG dynamics track both transient states and trait-like ability differences

## Introduction

Cognitive neuroscience seeks to identify the neural mechanisms underlying complex cognitive operations by relating measures of brain activity to behavioral performance. This approach has been enormously productive, but it relies largely on within-participant contrasts which compare neural activity across experimental conditions or correlate it with trial-level behavioral outcomes (Helfrich, Knight, & D’Esposito, in press). In the study of learning and memory, this logic takes the form of the subsequent memory paradigm, which compares brain activity recorded during encoding for items that are later recalled versus those that are not (Sanquist, Rohrbaugh, Syndulko, & Lindsley, 1980; Paller & Wagner, 2002). The resulting subsequent memory effect, or SME, has revealed characteristic patterns of spectral EEG activity that distinguish successful from unsuccessful encoding events (Rubinstein, Weidemann, Sperling, & Kahana, 2023; Weidemann & Kahana, 2021; Hanslmayr & Staudigl, 2014; Osipova et al., 2006). Yet, because these contrasts are computed within individuals, they reveal little about whether the same neural signatures also vary across individuals in ways that reflect stable differences in memory ability. The present work asks whether an electrophysiological signature that predicts memory within a person also differentiates people who consistently learn better than others.

Spectral EEG dynamics have been linked to the neural computations that support memory. In scalp EEG studies, low-theta (3–5 Hz) power over frontal channels typically increases during successful encoding, a pattern often interpreted as reflecting top-down control or contextual binding processes (Staudigl & Hanslmayr, 2013), though the direction of theta effects varies across recording modalities and referencing schemes (Herweg, Solomon, & Kahana, 2020). Gamma-band activity (30–100 Hz) has also been shown to increase during successful encoding across widespread hippocampo-cortical sites (Sederberg, Kahana, Howard, Donner, & Madsen, 2003; Burke, Ramayya, & Kahana, 2015), while alpha oscillations (8–14 Hz) are typically suppressed, a pattern associated with release from cortical inhibition and enhanced information processing (Jensen & Mazaheri, 2010; Hanslmayr, Staudigl, & Fellner, 2012). The joint expression of these effects, characterized by increased theta and gamma power (*T* ^+^,*G*^+^) coupled with decreased alpha (*A^−^*), has been termed the spectral T^+^A^-^G^+^ of successful memory (Katerman, Li, Pazdera, Keane, & Kahana, 2022; Broitman & Kahana, Submitted). However, Li, Pazdera, and Kahana (2024) demonstrated that when encoding classifiers control for the relation between serial position and recall probability, they rely primarily on alpha/beta decreases rather than the high-frequency activity increases present in classifiers trained without such controls. This finding suggests that previously reported scalp gamma SMEs partly reflected elevated gamma power at favorable serial positions rather than item-level encoding processes that operate independently of list position (see also Sederberg et al., 2006; Serruya, Sederberg, & Kahana, 2014; Kim, 2026). If neural activity systematically distinguishes successfully encoded items from forgotten ones within a person, a natural question is whether the magnitude of this distinction also varies across people in ways that reflect stable differences in mnemonic ability. This question arises particularly when one considers that healthy young adults performing under identical task conditions exhibit dramatic variation in recall performance (Underwood, Boruch, & Malmi, 1978; Healey, Crutchley, & Kahana, 2014), yet the source of these differences in encoding-related brain activity remains largely unexplored. While SME research has characterized the spectral signatures of encoding success at the group level in extensive detail, relatively few studies have examined whether SME magnitude correlates with individual differences in memory ability. Those that have done so come primarily from the fMRI literature, where Geissmann et al. (2023) showed in a sample of nearly 1,500 adults that hippocampal and prefrontal encoding differentiation positively predicted recall performance across individuals. A parallel line of fMRI work has linked hippocampal SME magnitude to associative memory performance across participants spanning young, middle-aged, and older adults (De Chastelaine et al., 2016). However, to our knowledge, no published electrophysiological study has systematically examined whether spectral SME magnitude covaries with individual differences in recall ability. This gap likely reflects practical barriers rather than lack of theoretical interest: individual-level SME estimates are noisy, requiring many trials per person to achieve adequate reliability (Ofen et al., 2021), and the sample sizes typical of EEG studies provide insufficient power to detect brain-behavior correlations across individuals (Elliott et al., 2020). Individual differences in spectral SMEs have been demonstrated before. Broitman, Healey, & Kahana, 2025 demonstrated that these effects weaken with chronological age, and suggested that this reduced T^+^A^-^G^+^ reflects degraded neural machinery. However, no studies have investigated whether this activity predicts individual memory ability among healthy young adults. Addressing this question requires a dataset that combines a large number of participants with sufficient measurement precision, a combination that requires multi-session, large-*N* datasets.

The present study takes a step towards incorporating neural measures in assessments of individual differences in memory function. We ask whether scalp EEG recordings taken during the learning phase of an experiment can be used to disentangle phasic fluctuations in memory-related neural signals from stable, trait-like neural markers of an individual’s mnemonic ability. To answer this question, we analyzed data from the Penn Electrophysiology of Encoding and Retrieval Study (PEERS; Kahana et al., 2024). This dataset consists of continuous high-density scalp EEG recordings taken as participants studied and subsequently recalled lists of items, and comprises more than 1.3 million encoding events across 98 individual participants. The scale and multi-session structure of this dataset provide sufficient measurement precision to estimate both within-participant variability in SMEs and stable between-participant differences in recall performance.

We focus on EEG activity within the theta, alpha, and gamma frequencies, because these frequency bands have been consistently implicated in scalp EEG studies of the SME (Rubinstein et al., 2023; Weidemann & Kahana, 2021; Hanslmayr & Staudigl, 2014; Osipova et al., 2006; Dougherty et al., in press). Previous work has shown that theta and gamma activity less strongly predict memory activity when participants divide attention across tasks during encoding, most likely because encoding-related resources get diverted to the secondary task (Broitman & Swallow, 2025; Broitman et al., 2025; Long & Kahana, 2016). These and other findings linking a reduced T^+^A^-^G^+^ effect to age-related memory deficits (Broitman et al., 2025) suggest that, under full attention, weaker memory performance should produce a weaker overall effects in healthy young adults. However, it remains unclear whether the magnitude of T^+^A^-^G^+^ varies among young adults who differ in their memory ability. If the SME reflects the efficient deployment of encoding resources, then better learners should show larger SMEs, because they more effectively engage the neural processes that support memory formation. If instead the SME reflects compensatory recruitment or greater fluctuation in encoding states, then better learners should show smaller SMEs, because their encoding processes operate more consistently across trials and require less phasic modulation to achieve successful learning. It is also possible that SME magnitude bears no systematic relationship to memory ability, if the neural processes it captures are equally available to good and poor learners alike. The present study tests these predictions by investigating whether the SME across theta, alpha, and gamma bands varies with individual differences in recall ability. We employ a mixed-effects linear modeling procedure which, like the method of Li et al. (2024), controls for serial position effects in the SME contrast, allowing us to determine which frequency components reflect encoding processes that operate across all list positions.

**Figure 1.**
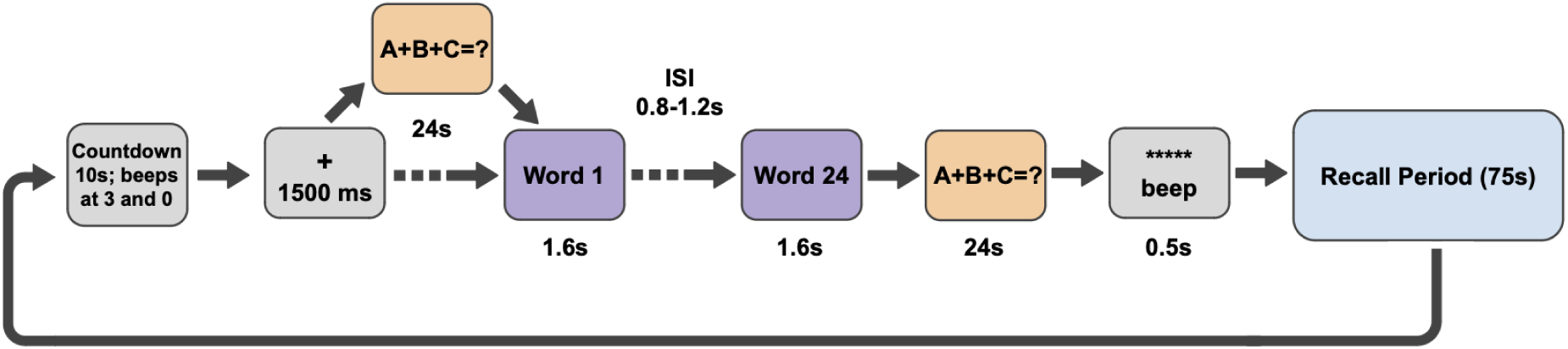
Schematic of PEERS methods. Participants studied lists of 24 words, and then vocally recalled the list items following a brief math distractor task.

## Methods

The data reported here come from the publicly available Penn Electrophysiology of Encoding and Retrieval Study (PEERS) described in Kahana et al. (2023). Participants in all PEERS experiments were recruited from among the students and staff at the University of Pennsylvania and neighboring institutions. This experiment tested delayed free recall of word lists (described as PEERS Experiment 4 in Kahana et al., 2023).

**Participants.** Participants were 98 young adults (52 female, 46 male), ranging from 18 to 30 years of age. The participant pool had a mean age of 21.45 (SD=3.06). Participants were recruited from the community surrounding the University of Pennsylvania. Each participant completed 24 sessions of multi-list delayed free recall.

**Experimental Task.** In each of the 24 experimental sessions, participants completed 24 lists of a delayed free recall task. In each list, participants first studied 24 session-unique English words. Words in each list were drawn without replacement from a pool of 576 common English words (see Kahana, Aggarwal, & Phan, 2018, for details regarding list construction). Each word appeared individually onscreen for 1,600 ms, and was followed by an interstimulus interval of 800–1,200 ms (uniformly distributed). Following the presentation of the last word, participants performed a distractor task for 24 s. The distractor task consisted of answering math problems of the form *A* + *B* + *C* = ?, where *A*, *B*, and *C* were positive, single-digit integers, though the answer could have been one or two digits. When a math problem was presented on the screen, participants typed the sum as quickly as possible. Participants were given a monetary bonus based on the speed and accuracy of their responses. After the post-encoding distractor task, there was a jittered delay of 1,200–1,400 ms, after which a tone sounded, a row of asterisks appeared, and the participant was given 75 s to freely recall the studied items. Participants were given a short break (about 3 min) after every eighth list in each session.

### EEG Preprocessing and Spectral Analysis

We recorded EEG with either a 129-channel EGI Geodesic Sensor Net in the Netstation acquisition environment, or with a 128-channel BioSemi Active Two system. We applied a 0.1 Hz high-pass filter to remove the EEG signals’ baseline drift over each session. In addition, we applied a fourth-order Butterworth notch filter with a 58-62 Hz stop-band to attenuate electrical line noise.

We also applied the same EEG preprocessing scheme as described in Katerman et. al (2022). We re-referenced recordings to the common average of all electrodes. When calculating the voltage mean across all electrodes, we excluded bad electrodes based on the channel variance and Hurst exponent. Specifically, raw session recordings were high-pass filtered at 0.5 Hz to reduce the impact of baseline drift on the variance and Hurst exponent. We then partitioned the EEG recording into three sections, separated by the two midsession breaks. Within each partition, we calculated the log-variance and Hurst exponent of each (non-electrooculogram) channel and Z-Scored across channels. We marked channels as bad if their Z-Scored log-variance exceeded +3 or fell below -3, or if their Z-Scored Hurst exponent exceeded 3 during any of the three partitions.

Because sampling rates varied between the two EEG systems, we resampled all scalp EEG data to a uniform 500 Hz. We then performed spectral decomposition using a Morlet wavelet transform (wave number = 6), implemented in Python using the Scipy package. Power was computed over a frequency range of 3–128 Hz. We spaced frequencies every 1 Hz in the range of 3–12 Hz, and every 4 Hz within 40–128 Hz, resulting in 33 frequencies of interest (Katerman et al., 2022). Our analyses focus on theta (3–5 Hz), alpha (10–12 Hz), and gamma (40–128 Hz) frequency bands. Spectral power was log-transformed and *z*-scored across trials within each frequency and electrode to normalize power estimates within each session.

### Event construction

We categorized each encoding event into one of two classes: successful memory events, defined as words that were subsequently recalled during the corresponding list’s recall period, and unsuccessful memory events, defined as words that were not recalled.

In accordance with prior work on neural SMEs (Long, Burke, & Kahana, 2014; Broitman et al., 2025), we extracted EEG data from 0 to 1,600 ms following each word onset. To minimize edge effects in spectral decomposition, we padded each EEG segment with a 400 ms buffer on both ends, resulting in a total window of -400 to 2,000 ms relative to word onset. We removed these buffer periods after computation.

### Computing the Subsequent Memory Effect

To quantify the relationship between neural activity during encoding and later memory performance, we computed the subsequent memory effect (SME) by contrasting spectral power for items that were subsequently recalled versus those that were not.

Spectral power estimates were computed for each encoding event across 33 principal frequencies (3–128 Hz), then collapsed across time (0-1,600 ms post-stimulus onset) and aggregated into three frequency bands: theta (3–5 Hz), alpha (10–12 Hz), and gamma (40–128 Hz). These bands were selected based on prior research linking frontal theta to associative binding and executive control, posterior alpha suppression to attentional gating, and posterior gamma to item-specific representations during successful encoding (Katerman et al., 2022; Wynn & Nyhus, 2022; Sederberg et al., 2007).

Power values were *z*-scored within each participant, frequency, and channel to normalize across sessions. We then collapsed spectral power across predefined scalp regions of interest (ROIs), guided by prior EEG studies using similar sensor arrays (Long & Kahana, 2014; Weidemann, Mollison, & Kahana, 2009). For scalp EEG, we initially defined eight ROIs partitioned into Left/Right, Anterior/Posterior and Superior/Inferior (Caudal) categories. These ROIs, as denoted in Fig. 2, were LAI, RAI, LAS, RAS, LPS, RPS, LPI, RPI.

Given prior evidence for bilateral expression of memory-related oscillatory effects (Katerman et al., 2022), we tested for hemispheric asymmetries by computing one-sample *t*-tests on the per-participant mean power differences between homologous left/right ROI pairs (e.g., LAI vs. RAI) across all three frequency bands. All twenty-four ROI-frequency pairs showed no significant hemispheric differences (*p ≥* 0.494), supporting bilateral consolidation of the signal.

Based on this empirical symmetry and prior literature emphasizing frontal theta and posterior alpha/gamma in memory encoding (Katerman et al., 2022; Wynn & Nyhus, 2022), we collapsed across hemispheres to define two primary ROIs for subsequent analyses: an anterior composite (LAI+RAI) and a posterior composite (LPS+RPS). We focus our primary analyses on anterior theta and posterior alpha and gamma.

### Statistical Modeling

This dataset comprised millions of encoding events hierarchically nested within sessions and participants. To account for this structure and properly separate within- and between-participant sources of variance, we constructed linear mixed-effects (LME) models using the lme4 package in R (Bates, Mächler, Bolker, & Walker, 2015). These trial-level models included fixed effects of subsequent memory (binary Recalled/Non-recalled labels). Depending on whether we were assessing session-level variance within participants or stable-trait like differences across participants, the models included additional fixed effects of session- or participant-level recall averages, and their interaction with trial-level recall. All models included random effects to account for participant-level differences in the effect of subsequent memory on spectral power, as well as random effects of serial position and list number to account for effects of the experimental structure on spectral power. All p-values reported below were FDR corrected for multiple comparisons.

**Participant Level Analyses.** To test whether spectral power reliably differentiated later-remembered from forgotten items at the group level, we fit a separate LME for each ROI-frequency band and experiment:

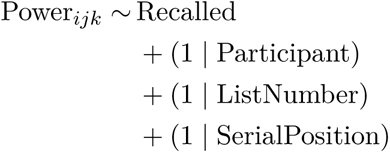

Where Power*_ijk_* denotes the specific spectral power at a given ROI-frequency pair for participant *i* at list position *j* and serial position *k*. We integrated session, list and serial position as mixed effects in order to control for confounding factors such as the primacy/recency effect. A significantly positive correlation indicates higher power during successful encoding (enhancement), whereas a negative correlation indicates suppression. These models provide a direct test of the canonical theta-alpha-gamma (T^+^A^-^G^+^) pattern: increased frontal theta and posterior gamma with concurrent posterior alpha suppression during successful encoding.

A likelihood ratio test confirmed that adding a random intercept for session did not improve model fit for any frequency band (all *χ*^2^(1) = 0, p = 1.000), so session was excluded from the final group-level model.

To assess whether the SME serves as a stable neural marker of individual recall ability, we fit a LME :

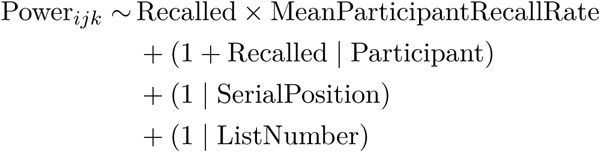

A significant interaction between Recall Status and Mean Participant Recall Rate would suggest that the SME correlates to a participant’s overall recall ability as a trait-like marker.

### Data and Code Sharing

All raw and pre-processed data, along with our analysis code, may be obtained from https://github.com/BenFalken/EEG-biomarkers-distinguish-good-and-poor-learners

## Results

As prior publications report detailed group-level behavioral results from this experiment (see Kahana et al., 2024), here we focus on the spectral SME and its relation to individual differences in overall recall rates. We report SMEs across the theta, alpha, and gamma frequencies, as well as their relation to individual recall performance. When analyzing the SME we control for serial position and list position effects using a mixed effects linear modeling procedure described in *Methods*. Serial position strongly influences both recall rates and spectral activity (e.g., Sederberg et al., 2006; Serruya et al., 2014), but not necessarily via the same channels. Controlling for serial position in the analysis of spectral SMEs helps to isolate factors related to putative endogenous factors that influence goodness of encoding (Li et al., 2024). Finally, we focus on specific ROIs based on previously published scalp EEG studies of the SME (Figure 2; Weidemann et al., 2009; Broitman et al., 2025; Long et al., 2014).

**Figure 2.**
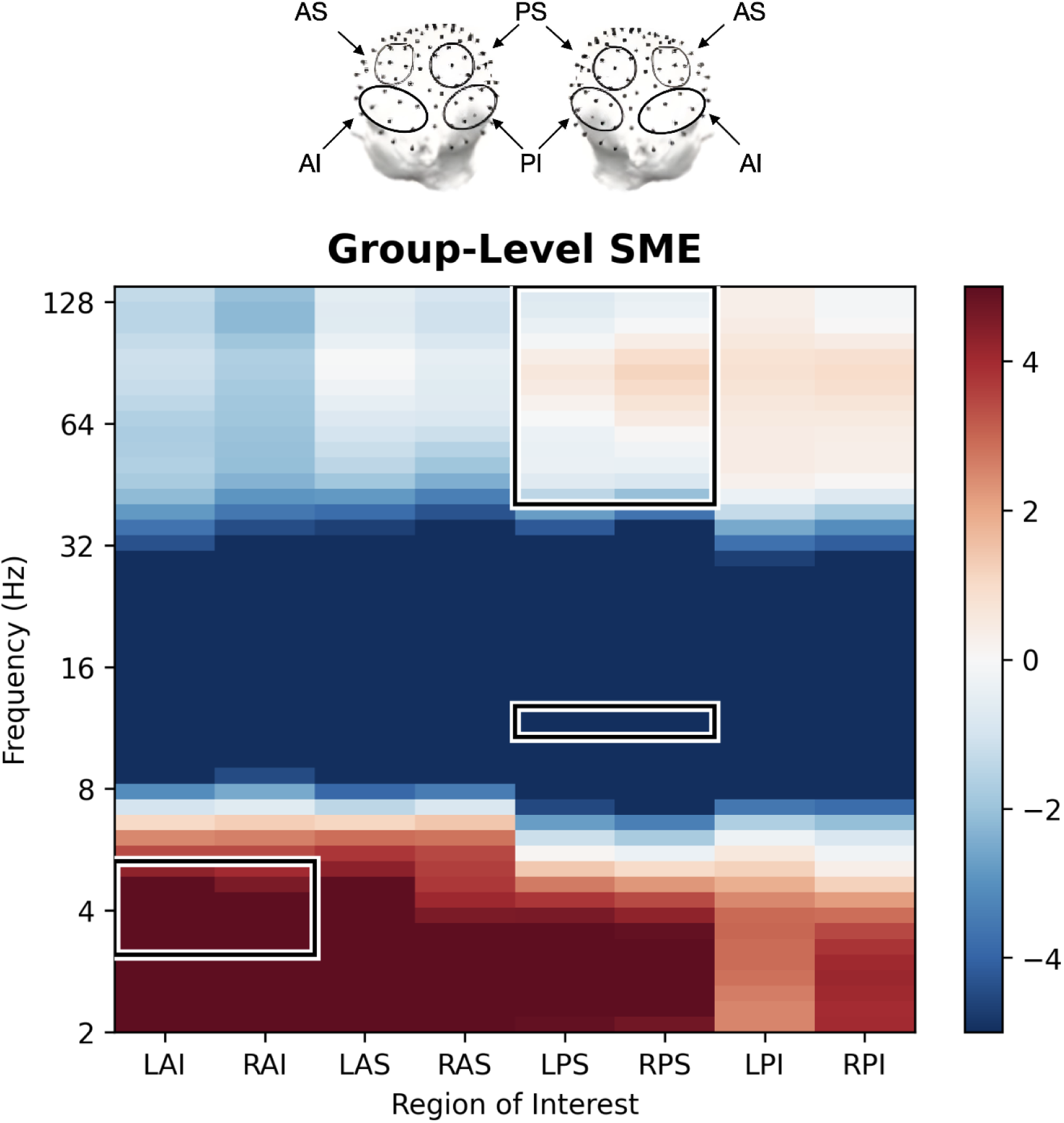
Above: The encircled electrodes comprise the eight principal ROIs, anterior/posterior and superior/inferior. When calculating the powers at each ROI, all electrodes are averaged into their respective groups. Below: Group-level SME calculated as the model-corrected average difference in z-scored log-power between recalled and non-recalled encoding events at every ROI-frequency pair, estimated via linear mixed-effects models controlling for participant, list, and serial position. Per-participant mean adjusted SME differences were then submitted to one-sample t-tests against zero, and the resulting t-statistics are displayed.

**Figure 3.**
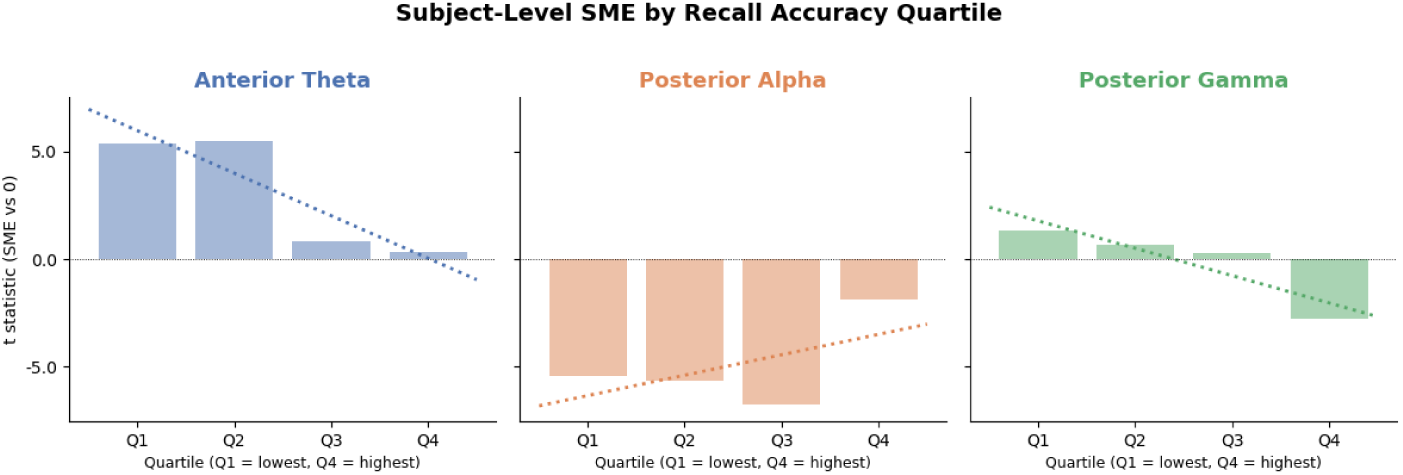
Participant-level SME was calculated by subtracting each participant’s, list’s, and serial position’s random intercept from trial-level power, then computing the mean difference in adjusted power between recalled and non-recalled events per participant. For each participant, a one-sample t-test of their adjusted recalled versus non-recalled power was computed against zero. Participant-level t-statistics were then aggregated by performance quartile (Q1 = lowest, Q4 = highest recall rate) and averaged within each bin. The dotted line denotes the line of best fit for the trait-level SME-recall trend.

**Group-Level Subsequent Memory Effects.** We first asked whether participants expressed subsequent memory effects in the theta, alpha, and gamma bands after controlling for list and serial position effects. ANOVAs on the model fits revealed a partial T^+^A^-^G^+^ profile: subsequent memory was associated with significant anterior theta enhancement (*β* = 0.0310 *±* 0.0010; F(1, 8975) = 1022.806; p < 0.001), coupled with posterior alpha suppression (*β* = -0.0665 *±* 0.0013; F(1, 55692) = 2455.494; p < 0.001). Gamma activity did not differentiate subsequently recalled from forgotten items (*β* = -0.0002 *±* 0.0012; F(1, 1259886) = 0.034; p = 0.854). This finding is consistent with prior work showing that associations between gamma power and subsequent memory primarily reflect primacy-related cognitive processes rather than endogenous states that facilitate encoding (Sederberg et al., 2006; Serruya et al., 2014; Li et al., 2024).

**Does the SME predict participant-level recall?** Figure 3 presents the SME magnitude across individual recall performance quartiles. Across all three component frequencies, better participant-level memory was predicted by a weakened T^+^A^-^G^+^ SME. Among higher-performing individuals, there was a weaker association between subsequent memory and positive theta (*β* = -0.1235 *±* 0.0283; F(1,97.1)=19.123; p<0.001), negative alpha (*β* = 0.0857 *±* 0.0342; F(1,96.5)=6.263; p=0.014), and positive gamma (*β* = -0.0499 *±* 0.0231; F(1,94.5)=4.671; p=0.033).

**Post-hoc analysis: Do better learners display less neural variability?** A possible explanation for the weakened SMEs among better learners is that these individuals display reduced trial-to-trial variability in encoding-related neural activity. We therefore investigated whether participant-level recall ability correlated with across-trial neural variability within each of the three component frequencies of T^+^A^-^G^+^ . To test for this possibility, we first computed log-power at each encoding trial within each frequency and electrode. We used the linear mixed-effects models described above to adjust each logpower value for random effects of participant, list, and serial position. We then computed the standard deviation of the model-adjusted log-power values for each participant within each frequency band and ROI. For each frequency band, we then computed a Pearson correlation between each participant’s across-trial standard deviation of log-power and their average recall probability.

**Figure 4.**
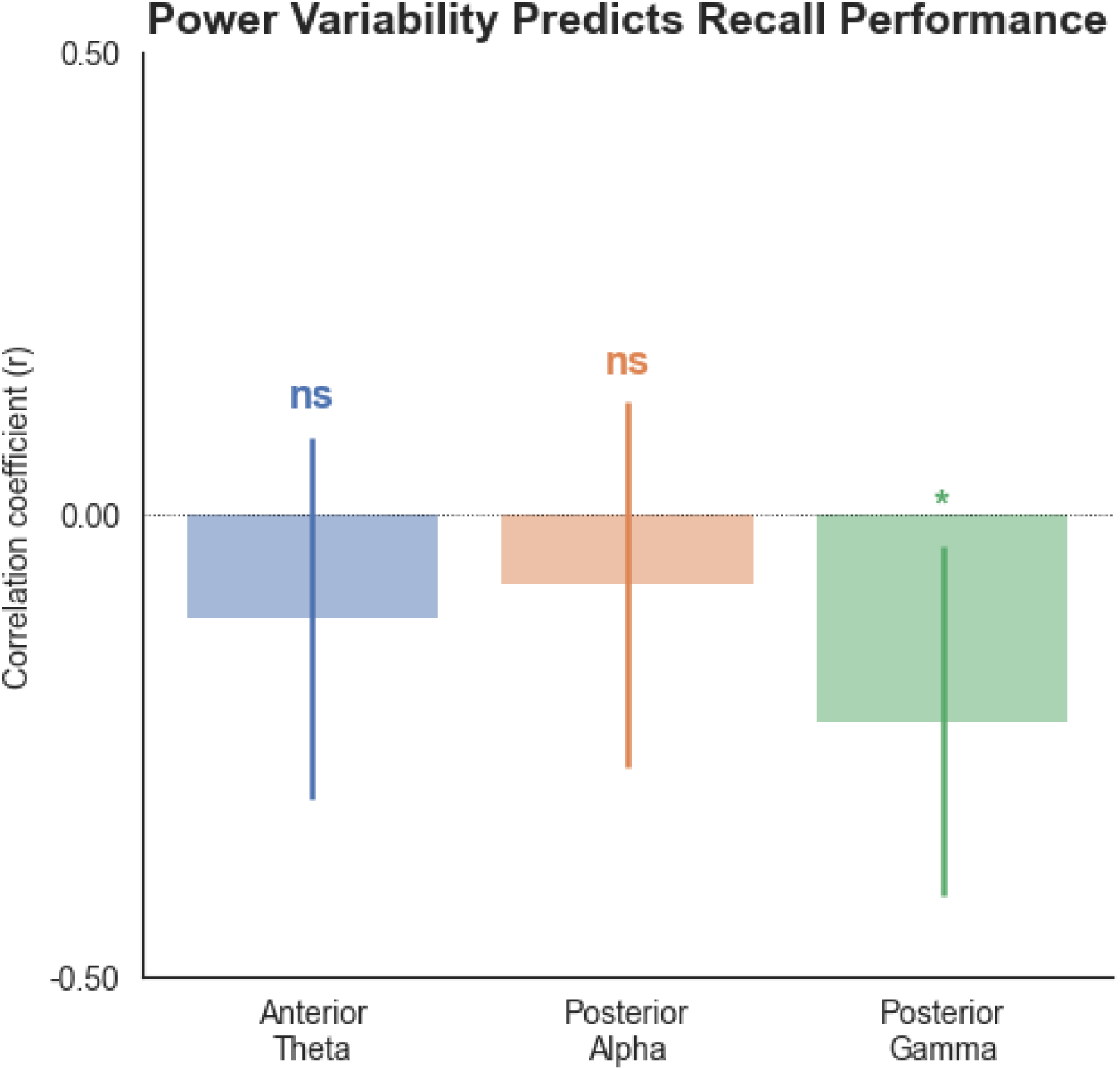
Relationship between subject-level neural variability and recall performance across the three component frequencies of T^+^A^-^G^+^ . For each participant, we computed the standard deviation of model-adjusted log-power across all encoding events, separately for anterior theta, posterior alpha, and posterior gamma. We then computed the Pearson correlation between each participant’s standard deviation and their average recall probability. Bars denote the correlation coefficient (r) for each frequency band; error bars denote SEM. Asterisks denote significance following FDR correction.

Across all three component frequencies, greater participant-level recall probability was associated with lower neural variability across encoding trials (Figure 4). Following FDR correction, this correlation was non-significant across the theta (*r* = *−*0.112*, p* = .273) and alpha (*r* = *−*0.148*, p* = .147) frequencies. However, the correlation was significant within the gamma band (*r* = *−*0.223*, p* = .027). Though these effects are modest and should be interpreted with caution, they are consistent with the possibility that the reduced SMEs across higher-performing individuals stems from reduced levels of trial-to-trial neural variability.

## Discussion

Spectral SMEs such as the T^+^A^-^G^+^ pattern reliably distinguish subsequently remembered from forgotten items. The present study asked whether this within-person neural contrast also varies across individuals in ways that reflect stable differences in mnemonic ability. After controlling for serial position and list effects using mixed-effects models, we found increased theta and decreased alpha activity predicted successful encoding, while gamma power did not demonstrate any reliable effects. However, SME magnitude in all three component frequencies varied across individuals. Better learners exhibited attenuated SMEs across all three frequency bands, indicating that the same neural dynamics that mark successful encoding on a given trial are weaker overall in the people who remember the most. This challenges the assumption that larger SMEs reflect more effective encoding, and suggests that T^+^A^-^G^+^ magnitude captures stable, trait-like differences in learning efficiency.

The absence of a reliable gamma SME after controlling for serial position and list effects is consistent with growing evidence that scalp gamma associations with subsequent memory largely reflect primacy-related processes rather than item-level encoding mechanisms that operate across all list positions (Sederberg et al., 2006; Serruya et al., 2014; Li et al., 2024). Our mixed-effects models, which included serial position as a random effect, support this conclusion. After accounting for positional variance, gamma does not reliably differentiate recalled from forgotten items at the group level. Alpha suppression, by contrast, was robust to these controls and produced the strongest group-level SME, while theta positivity also remained significant. This dissociation adds to the evidence that the components of T^+^A^-^G^+^ do not behave as a unified signal, and that alpha and theta SMEs may be less susceptible to serial position confounds than gamma-band activity.

Better learners exhibited weaker SMEs across all three frequency bands, meaning the neural contrast between recalled and forgotten items was most pronounced among those who remembered the least. We considered the possibility that better learners maintain more stable encoding states across trials, reducing the difference in neural activity between recalled and forgotten items. A post-hoc analysis found that neural variability was negatively correlated with recall performance across all three frequency bands, though only the gamma-band correlation was statistically significant after correction. This pattern suggests that poorer learners may fluctuate more between neural states that do and do not support successful encoding, producing both lower recall rates and larger contrasts between recalled and forgotten items. Better learners, by contrast, may operate within a narrower and more consistently effective range of encoding states, leaving less room for the recalled/forgotten contrast to emerge.

Despite evidence that T^+^A^-^G^+^ reflects multiple frequency-specific cognitive processes (Broitman & Kahana, 2026; Fellner et al., 2019), better learners exhibited reduced SMEs across all three component frequencies, suggesting a common upstream cause. Greater theta SMEs in poorer learners may reflect increased reliance on effortful, controlled encoding processes. Frontal theta has been linked to top-down control and context binding (Herweg et al., 2020; Staudigl & Hanslmayr, 2013), and poorer learners may need to recruit these processes more heavily to successfully encode an item. Greater alpha suppression may reflect the need for greater effort to reach an engaged cortical state that better learners sustain by default. According to the gating-by-inhibition framework (Jensen & Mazaheri, 2010), alpha oscillations reflect the suppression of task-irrelevant information. If better learners display higher baseline alpha activity (Klimesch, 1999), this could reflect more effective default filtering that reduces the need for large phasic adjustments during encoding. The between-participant gamma effect, though significant, should be interpreted with caution given the absence of a reliable group-level gamma SME after serial position controls. One possibility is that it reflects reduced effortful neural computation among better learners, consistent with broadband gamma’s proposed role as an index of local cortical processing demand (Ray & Maunsell, 2011). An additional possibility is that the gamma SME reverses among better learners. In this case, gamma activity may disrupt task-oriented states that are beneficial to encoding (Broitman & Swallow, 2025).

Our results contrast with those reported by Broitman et al. (2025), who found that a weakened T^+^A^-^G^+^ SME reflected age-related memory impairments. We must therefore consider multiple reasons for why SMEs can be weak. One possibility is that age-related memory impairments stem from increased interference between study and retrieval (Craik & Jennings, 1992; Naveh-Benjamin, 2000; Kahana, Dolan, Sauder, & Wingfield, 2005), in which case even a clear neural signal of good encoding would have less predictive value for which items get retrieved later. Among high-performing young adults, weakened SMEs might instead reflect a reduced dependence on achieving a specific neural profile for successful encoding. If both interpretations are correct, then the same reduction in SME magnitude would have opposite functional implications depending on the population being studied.

### Study Limitations

Several methodological limitations might reduce the inferential reach of our findings. Our dataset primarily consisted of young, highly educated participants from a university community. This homogeneity might limit the generalizability of our conclusions, and future studies should seek to recruit populations with greater variability in age and educational background. The subsequent memory paradigm also introduces a structural confound. Better performing participants contribute more recalled items to the SME contrast, so differences in SME magnitude could partly reflect differences in the ratio of successful to unsuccessful events rather than purely mechanistic differences in encoding. Future work could address this by using continuous neural predictors of memory, such as regularized logistic regression, rather than relying on post-hoc contrasts between recalled and forgotten items.

## Conclusions

By combining large-scale EEG with hierarchical mixed-effects modeling, the present study demonstrates that spectral SMEs vary across individuals in ways that reflect stable differences in learning ability. This approach reveals that the same spectral dynamics that predict memory on a given trial also vary systematically across individuals in ways that are undetectable using standard within-participant analyses. An important next step is to examine whether differences in T^+^A^-^G^+^ across individuals also emerge at retrieval. If high-performing young adults exhibit preserved T^+^A^-^G^+^ effects during successful recall despite weakened encoding SMEs, this would suggest that the weakened encoding contrast reflects reduced dependence on phasic spectral activity rather than reduced engagement of the memory system. More broadly, the ability to assess stable individual differences in cognitive ability from spectral EEG has translational potential. If trait-level markers of encoding efficiency can be reliably extracted from the EEG signal, they could inform the early detection of memory disorders, and the development of individualized therapeutic interventions that adapt to a person’s neural profile.

